# Elevation of linkage disequilibrium above neutral expectations in ancestral and derived populations of *Drosophila melanogaster*

**DOI:** 10.1101/029942

**Authors:** Nandita R. Garud, Dmitri A. Petrov

## Abstract

The extent to which selection and demography impact patterns of genetic diversity in natural populations of *Drosophila melanogaster* is yet to be fully understood. We previously observed that the pattern of LD at scales of ~10 kb in the Drosophila Genetic Reference Panel (DGRP), consisting of 145 inbred strains from Raleigh, North Carolina, measured both between pairs of sites and as haplotype homozygosity, is elevated above neutral demographic expectations. Further, we demonstrated that signatures of strong and recent soft sweeps are abundant. However, the extent to which this pattern is specific to this derived and admixed population is unknown. Neither is it clear whether such a pattern may have arisen as a consequence of the extensive inbreeding performed to generate the DGRP data. Here we analyze > 100 fully sequenced strains from Zambia, an ancestral population to the Raleigh population, that has experienced little to no admixture and was generated by sequencing haploid embryos rather than inbred strains. This data set allows us to determine whether patterns of elevated LD and signatures of abundant soft sweeps are generic to multiple populations of *D. melanogaster* or whether they are generated either by inbreeding, bottlenecks or admixture in the DGRP dataset. We find an elevation in long-range LD and haplotype homozygosity in the Zambian dataset, confirming the result from the DGRP data set. This elevation in LD and haplotype structure remains even after controlling for many sources of LD in the data including genomic inversions, admixture, population substructure, close relatedness of individual strains, and recombination rate variation. Furthermore, signatures of partial soft sweeps similar to those found in the DGRP are common in Zambia. These results suggest that while the selective forces and sources of adaptive mutations may differ in Zambia and Raleigh, elevated long-range LD and signatures of soft sweeps are generic in *D. melanogaster*.

## INTRODUCTION

Disentangling the effects of demography and selection on patterns of genomic variation remains a central challenge in evolutionary biology. Until recently, inference of demography and selection relied primarily on short genomic fragments sampled from a limited number of individuals (e.g. Duchen *et al*. 2013; Gutenkunst *et al*. 2009; Li and Stephan 2006; Molina *et al*. 2001; Pritchard *et al*. 2000; Thornton and Andolfatto 2006). While such data provided important insights into the evolutionary forces acting on populations, they were ultimately limited by both sample size and the physical scale in which patterns of polymorphism and linkage could be investigated. The recent availability of whole genome sequences from multiple individuals with populations in a variety of species (e.g Abecasis *et al*. 2010; Cao *et al*. 2011; Mackay *et al*. 2012) is enabling us to examine long-range patterns of LD (~10Kb), measured either as correlations between pairs of sites or as haplotype homozygosity. LD offers powerful insights into selective and demographic processes shaping genetic variation in a natural population, such as bottleneck or admixture demographic forces (Pennings and Hermisson 2006; Pokalyuk 2012; Wall and Pritchard 2003; Andolfatto and Przeworski 2000). In addition, elevated haplotype homozygosity is very sensitive to recent and strong adaptation leading to both classical hard and soft selective sweeps (Garud *et al*. 2015; Garud and Rosenberg 2015; Pennings and Hermisson 2006).

In the case of *Drosophila melanogaster,* the most recent demographic models were inferred using short intergenic and intronic sequence fragments on the X chromosome of lengths no longer than a few hundred base pairs in tens of individuals (Duchen *et al*. 2013). Because the data were so sparse, models assuming independence between polymorphic sites were fit to statistics sensitive to the site frequency spectrum as well as to short-scale linkage disequilibrium measuring the correlation between polymorphic sites separated by short distances (Duchen *et al*. 2013).

The availability of the Drosophila Genetic Reference Panel (DGRP), consisting of 162 fully sequenced *D. melanogaster* strains from a single population in Raleigh, North carolina (Mackay *et al*. 2012), offered the opportunity to study genome-wide signatures of demography and selection in a deep sample. Recently, we analyzed patterns of LD and haplotype homozygosity at distances of ~10Kb in the DGRP data set (Garud *et al*. 2015) and compared these patterns with neutral expectations generated under several demographic models of North American *D. melanogaster,* including two constant *N_e_* models, two bottleneck models, and two recently inferred admixture models (Duchen *et al*. 2013), one of which is considered to be the most accurate model of the admixture between European and African flies founding North American *D. melanogaster*. We found that LD and haplotype homozygosity in the DGRP data is elevated relative to expectations under any of the demographic models tested and that this elevation remained even after controlling for several potential sources of elevated LD including close relatedness between individuals, presence of genomic inversions, recombination rate variation, and population substructure.

In addition, we identified regions in the DGRP data with the elevated haplotype homozygosity, measured using statistic H12, designed to detect both hard and soft sweeps Garud *et al*. 2015. H12 calculates haplotype homozygosity after combining the frequencies of the first and second most common haplotypes in order to increase the power to detect soft sweeps. Among the three highest-ranking candidates in the H12 scan were the prominent known cases of soft sweeps at the loci *Ace, Cyp6g1,* and *CHKov1* (Karasov *et al*. 2010; Magwire *et al*. 2011; Schmidt *et al*. 2010). All 50 outlier H12 peaks in the DGRP data showed signatures of partial soft sweeps, suggesting that soft sweeps were common in the DGRP data (but see Schrider *et al*. 2015).

The extent to which soft sweeps are specific to this Raleigh population remains unknown. Several factors challenge the interpretation of the results of the DGRP data set and we seek to address them in this paper. First, the Raleigh population, like all other North American populations of *D. melanogaster,* is a recently derived population resulting from admixture between European and African lineages of flies (Bergland *et al*. 2015; Duchen *et al*. 2013; Pool *et al*. 2012). The effects of recent admixture on patterns of LD and haplotype homozygosity are still yet to be fully understood, and furthermore the true demographic history of North American flies is undoubtedly much more complex than our current models can possibly capture. For example, Pool 2015 showed that epistatic interactions between African alleles and European alleles are abundant in the North American population and that these combinations of alleles appear to be experiencing selection against them.

Second, the flies comprising the DGRP data set were extensively inbred to result in mostly homozygous genomes. This inbreeding process left behind tracts of residual heterozygosity which were suggested by Houle and Marquez 2015 to result from epistatic interactions between inversions (particularly heterozygous inversions that may be lethal as homozygotes) maintained on different chromosomes or other deleterious mutations in repulsion. Unfortunately this may result in elevations in long-range linkage disequilibrium since regions of the genome bearing deleterious mutations may have be to maintained in-sync with other potentially non-deleterious regions of the genome, otherwise the compound effect of two deleterious regions present in the genome could be lethal.

Recently, the Drosophila Genome Nexus (DGN) dataset was published (Lack *et al*. 2015), consisting of 205 DGRP strains from Raleigh (inclusive of the original 162 DGRP strains available in the initial release) as well as 197 strains from Zambia. The Zambian data set offers the opportunity to test whether signatures observed in the Raleigh population are generic to multiple populations of *D. melanogaster*. Importantly, the two features of extensive admixture in the Raleigh population and inbreeding of the flies to generate the data set are absent from the Zambian lines. First, as the likely ancestral *D. melanogaster* population, the Zambian population has experienced relatively little admixture (Pool *et al*. 2012). Second, since Zambian flies have a much higher level of genetic diversity (Pool *et al*. 2012) and deleterious load, the flies cannot be fully inbred to generate phased haplotypes for the sequenced data set. Instead, Langley *et al*. 2011 used a crossing technique to generate and sequence haploid embryos from isofemale strains. The lack of admixture and lower levels of inbreeding in the Zambian lines set make a comparison of patterns of LD and haplotype homozygosity between the Zambian data and the Raleigh data particularly informative in understanding general patterns of soft selective sweeps in *D. melanogaster*.

In this paper, we compare patterns of linkage disequilibrium and haplotype homozygosity in flies sampled from Raleigh and Zambia. We find that LD and haplotype homozygosity in both Raleigh and Zambian lines are elevated relative to neutral expectations, suggesting that elevated LD and haplotype homozygosity is a generic feature of multiple *D. melanogaster* populations. Furthermore, our H12 scan of the Zambian lines reveals several peaks in genomic locations different from that of Raleigh, many of which show signatures of soft sweeps with multiple haplotypes at high frequencies. In the Zambian data set, we recover peaks at two positive controls, *Ace* and *Cyp6g1,* but these are not the highest-ranking peaks as in the Raleigh data set, reflecting the lower frequencies at which the adaptive mutations are found in ancestral, African populations than in derived populations. We do not recover a peak at *CHKov1*, which is a negative control in African populations since the adaptive mutation has been shown to be present at high frequencies only in non-African populations (Aminetzach *et al*. 2005). While the patterns of elevated long-range LD and multiple haplotypes at high frequencies in both populations show that soft sweeps may be common in both populations, the differences in genomic locations of individual selective sweeps is consistent with different underlying selective pressures and evolutionary processes in the two populations.

## RESULTS

### Data

In this paper we contrast levels of LD and haplotype homozygosity in the Raleigh and Zambian populations. In our analysis, we masked previously inferred regions of identity by descent (IBD), genomic inversions, and tracts of residual heterozygosity resulting from the inbreeding of the Raleigh strains (Corbett-Detig and Hartl 2012; Huang *et al*. 2014; Lack *et al*. 2015). For Zambian strains, we also masked previously inferred regions of admixture from Europe into Africa (Lack *et al*. 2015; Pool *et al*. 2012) (Methods). Finally, we excluded regions of the genome with low recombination rates < 1 centimorgan/megabase (cM/Mb) (Comeron *et al*. 2012). The extensive masking of this data set resulted in large tracts of missing data across several individuals and introduced variation in the sample size at different sites in the genome. To account for this variation, we sub-sampled both the Raleigh and Zambian data to 100 strains, excluding the worst strains in terms of missing data (Methods).

Later in this paper we perform a genome-wide H12 scan of the Zambian and Raleigh data sets to identify individual selective events in each population. To maximize our coverage and power to identify selective sweeps, we applied less stringent filters to the data by masking tracts of residual heterozygosity and additional heterozygous sites as before, but not masking any inversions, regions of inferred admixture, or IBD tracts. Instead, we *a posteriori* tested for enrichment of selective sweeps in genomic regions with inversions present at high frequency in the data set. In addition, instead of masking IBD tracts, we excluded all individuals with genome-wide IBD levels with at least one other individual at > 20% (Methods). This resulted in 178 and 188 strains in the Raleigh and Zambian data sets, respectively. We further down-sampled the data sets to 145 strains excluding the strains with the highest amount of missing data (Methods) to match the sample depth of our previous H12 scan (Garud *et al*. 2015). Finally, we excluded regions of the genome with recombination rates < 0.5 cM/Mb (Comeron *et al*. 2012). We refer to this second data set as the less stringently filtered data set.

### Slow decay of linkage disequilibrium in the Zambian and Raleigh populations

In Figure 1a, we compare the decay in LD in the Zambian data with expectations under neutral demographic models fit to the data. We inferred four simple neutral demographic models fit to short introns in the Zambian lines using the program DaDi (Gutenkunst *et al*. 2009): a constant *N_e_* demographic model, a bottleneck model, a growth model, and a model of a bottleneck followed by expansion (hereafter, ‘bottlegrowth’) (Methods). *S* and π for each model match empirical estimates from short-intron data in the Zambian population (Table 1). The log-likelihoods of each model and their inferred parameters are presented in Table 2. it is clear based on the log likelihoods that the bottlegrowth model and bottleneck models have the best fits. However since even the bottleneck model experiences a growth in population size, we decided to focus on the bottlegrowth model. In addition, we estimated the decay in LD in simulations of the constant *N_e_* and growth models as points of comparison. Since low recombination can result in elevated LD, we excluded regions of the genome with recombination rates < 1 cM/Mb (Comeron *et al*. 2012) and performed all neutral simulations with this recombination rate. This recombination rate should be conservative for our purposes as this is the lowest recombination rate in our data.

**Table 1:** *S* and *π* measured in neutral demographic models of Zambian and Raleigh populations of *D. melanogaster*. (A) *S* and *π* estimates for Zambian short intron data and demographic models fit to Zambian short intron data. (B) *S* and *π* estimates for Raleigh short intron data and demographic models fit to Raleigh short intron data. Estimates of *S* and *π* were averaged over 30,000 simulations of 10,000 bps for each demographic model.

(A) Zambia
| Data/model | $S/\text{bp}$ | $\pi/\text{bp}$ |
| --- | --- | --- |
| ZI short intron data | 10.1% | 1.6% |
| Constant $N_e=4.8 \times 10^6$ model | 9.9% | 1.9% |
| Growth model | 10.2% | 1.1% |
| Bottlegrowth model | 12.1% | 1.7% |

(B) Raleigh
| Data/model | $S/\text{bp}$ | $\pi/\text{bp}$ |
| --- | --- | --- |
| RA short intron data | 5.8% | 1.2% |
| Admixture model | 5.8% | 1.1% |
| Admixture + bottleneck model | 5.6% | 1.3% |
| Constant $N_e=10^6$ model | 2.3% | 0.4% |
| Constant $N_e=2.7 \times 10^6$ model | 5.8% | 1.1% |
| Severe short bottleneck model | 5.7% | 1.1% |
| Shallow long bottleneck model | 5.5% | 1.1% |

**Figure 1:**
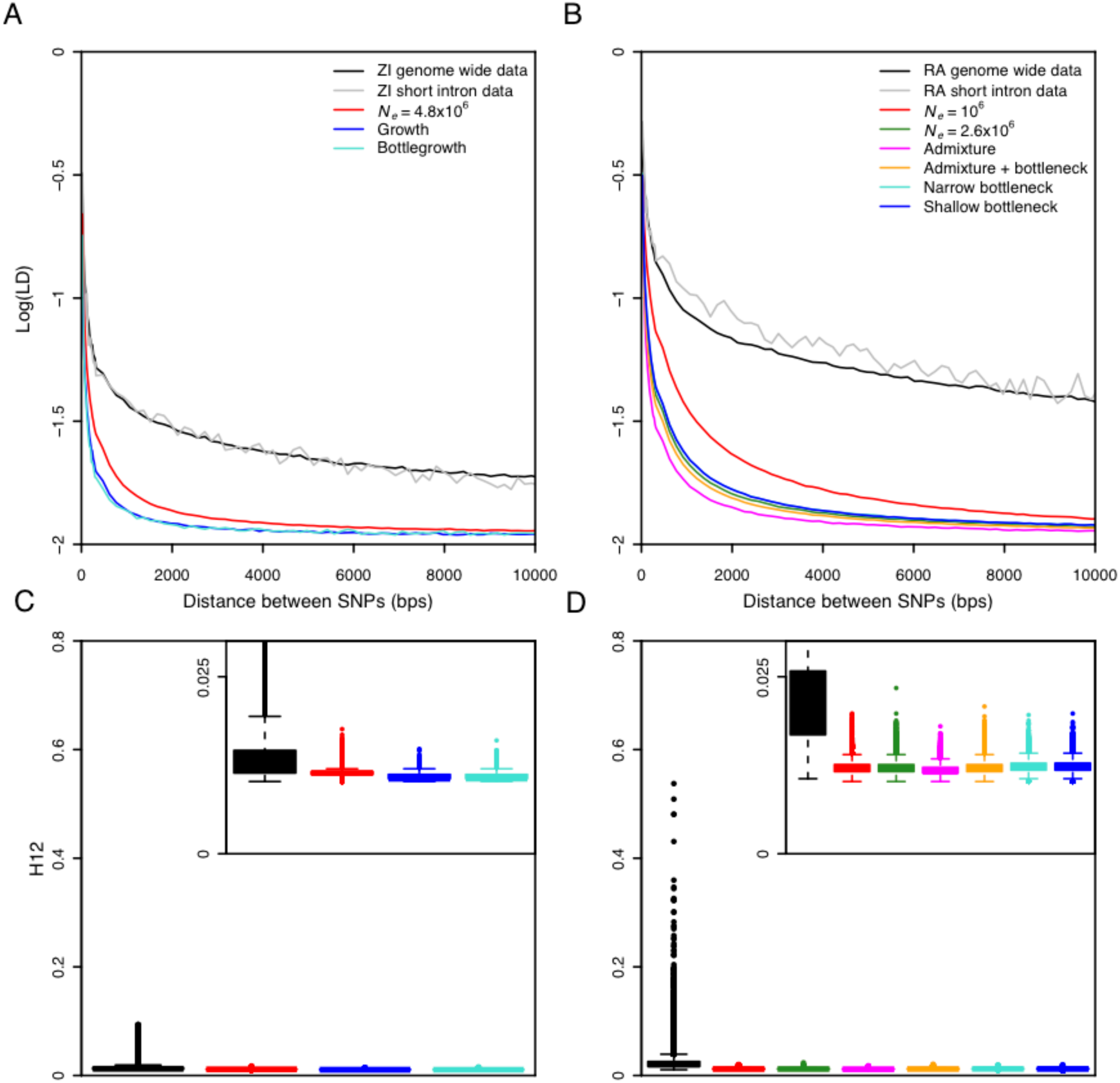
Elevated long-range LD and haplotype homozygosity in Zambia and Raleigh populations. LD in Zambia (ZI) (A) and Raleigh (RA) (B) is elevated as compared to any neutral demographic model, especially for long distances. pairwise LD was calculated in Zambia and Raleigh data for regions of the *D. melanogaster* genome with *ϱ ≥* 10^−6^ cM/bp. Neutral demographic simulations were generated with *ϱ* = 10^−6^ cM/bp. Pairwise LD was averaged over 10^7^ simulations in each neutral demographic scenario. Genome-wide H12 values measured in 801 SNP windows in Zambia data (C) and 401 SNP windows in Raleigh data (D) are elevated as compared to expectations under any neutral demographic model tested. H12 values were measured in sample sizes of 100 and in genomic regions with *ϱ* ≥ 10^−6^ cM/bp. H12 values were measured in neutral demographic simulations of 100 individuals generated with *ϱ* = 10^−6^ cM/bp. Plotted are the result of approximately 1.5×10^5^ simulations under each neutral demographic model, representing ten times the number of analysis windows in the data.

We observe a substantial elevation in LD in the data relative to neutral expectations under all demographic models (Figure 1a). We estimated LD for different minor allele frequency (MAF) classes of SNPs (Methods) where both SNPs in an LD pair had MAFs between 0.05 and 0.5 (Figure 1a), 0.3 and 0.5 (Figure S1a), and 0 and 0.05 (Figure S1b). Using a two-sided Wilcoxon rank sum test to test the significance of the deviation in LD from neutral expectations, the *P*-value was <10^−15^ in all cases. The deviation in LD in the data from expectations is apparent at short distances less than 100 basepairs, consistent with observations of Przeworski *et al*. 2001 for multiple species of Drosophila, as well as for long distances, consistent with our previous results (Garud *et al*. 2015).

In Figures 1b, S1c and S1d, we reiterate our previous result in Garud *et al*. 2015 in the Raleigh data set and confirm that there is an elevation in LD in the Raleigh population relative to the six neutral demographic models fit to the Raleigh data in terms of *S* and *π* (Table 1). In contrast to the LD in the Zambian data set, the deviation in LD from neutral expectations is even more exaggerated in the Raleigh data set (For all MAFs conditioned on and demographic models tested, the *P*-value is < 10^−15^ for a two-sided Wilcoxon rank sum test). The elevation of LD in both Zambian and Raleigh data suggest that elevated LD is a generic feature of multiple *D. melanogaster* populations.

To confirm that the elevation in LD is not driven by any anomaly in the data such as unaccounted related individuals, inversions, or strains with unusually high polymorphism or sequencing error rates, we subsampled the Zambian and Raleigh data sets 10 times to 50 strains to test whether any subset of individuals may be driving the elevation in LD. In all subsampled data sets, LD remains elevated in contrast to expected LD measured in all tested demographic models simulated with a sample size of 50 (*P*-value < 10^−15^ for all scenarios tested, two-sided Wilcoxon rank sum test) (Figure S2) suggesting that the elevation in LD is driven by population-wide phenomena.

### Elevation of haplotype homozygosity in the Zambian and Raleigh populations

We compared the distribution of haplotype homozygosity measured with H12 in the Zambian and Raleigh data sets with the distribution of H12 values measured in neutral demographic scenarios fit to the two populations (Figures 1c and 1d). H12 is designed to detect both hard and soft sweeps and is calculated by combining the frequencies of the two most abundant haplotypes in an analysis window (Garud *et al*. 2015). For the Raleigh data, we applied the statistic in analysis windows of 401 SNPs (200 SNPs on each side of the central SNP of each window) with the centers of each window iterated by 50 SNPs. 401 SNPs, corresponds to ~10Kb in the Raleigh data (Garud *et al*. 2015). In the Zambian data, however, 801 SNPs corresponds to approximately ~10Kb. Therefore, we applied the statistic in 801 SNP windows randomly down sampled once to 401 SNPs in the Zambian data to have comparable analysis windows sizes in terms of SNPs and base pairs for both scans. A window of this length scale is unlikely to generate high values of LD or haplotype by chance under neutrality but instead should have high power to identify significantly elevated haplotype homozygosity arising from strong selection events with *s* = 0.05% or greater (Garud *et al*. 2015). In our application of the statistic, we excluded regions of low recombination (<1 cM/Mb) (Comeron *et al*. 2012) because low levels of recombination can generate high homozygosity. We performed 150,000 simulations of each neutral demographic scenario, representing approximately 10 times the number of analysis windows observed in the data.

Figures 1c and 1d show that there is a long tail of high H12 values in both the Zambian and Raleigh data sets relative to the distribution of H12 measured in simulations under all neutral demographic models tested, confirming our previous results (Garud *et al*. 2015) showing that all the neutral demographic scenarios we tested are unable to generate the haplotype structure observed in the Raleigh data by chance (*P*-value < 10^−15^, Wilcoxon rank sum test, for all neutral demographic scenarios tested). The elevation in H12 values in the tail is muted in the Zambian data set with the highest H12 value reaching only 0.0936, whereas in the Raleigh data set the highest value reaches 0.5374. In the Raleigh data set, the median H12 value (0.0202) is sharply elevated above the median H12 value measured in simulations (~0.012 across all demographic scenarios) and ~68% of the windows in the data have H12 values greater than the 1-per-genome false discovery rate values corresponding to the 10^th^ most extreme H12 value in the distributions of H12 values measured in simulations. In the Zambian data set, we found that the elevation of the median H12 value in the data (0.0124) is also extreme relative to the median H12 value measured in simulations (~0.0112 across all demographic scenarios) with ~26% of the windows in the data having H12 values above the 1-per-genome false discovery rate.

In the next section we perform a second genome-wide H12 scan to identify individual selective sweeps in less stringently masked data sets where only tracts of residual heterozygosity and any additional remaining heterozygous sites are masked. This less stringently masked data set provides us greater power and coverage to identify selective sweeps. To measure the effect of the extensive masking of regions of admixture, IBD, inversions, and low recombination on the distribution of H12 values, we compared the distributions of H12 values measured in the stringently and less-stringently masked data sets finding that the distributions are similar (Figure S3, Figures 1c and 1d), with an elevation of the tails and medians of H12 values in the data relative to neutral demographic simulations.

### Identification of selective sweeps in the Zambian and Raleigh data

We performed a genome-wide H12 scan to identify individual selective sweeps in the Zambian and Raleigh data in the less stringently filtered data sets (Figure 2). For the Raleigh data, we applied the statistic in windows of 401 SNPs with the window centers separated by 50 SNPs. For the Zambian data, we applied the statistic in 801 SNP windows randomly down sampled once to 401 SNPs. To test the effect of our window size choice on the locations of H12 peaks in the Zambian data, we applied H12 in windows of 801 SNPs in the Zambian data in Figure S4 and found that our results do not change substantially. In our analyses, we excluded genomic regions with low recombination rates < 0.5 cM/Mb (Comeron *et al*. 2012) because low recombination rates can result in high homozygosity values.

**Figure 2:**
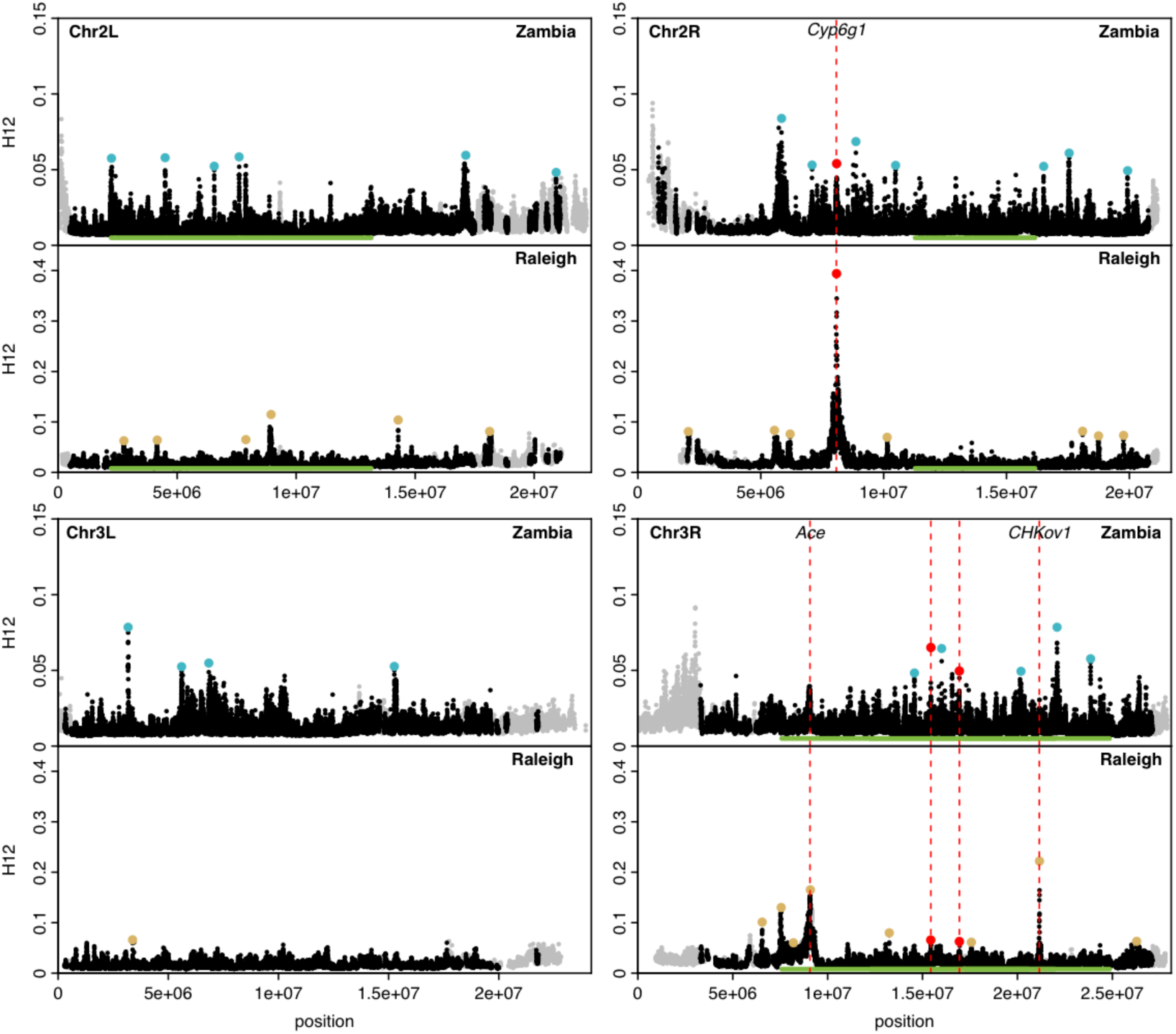
H12 scan in Zambian and Raleigh data. We performed an H12 scan in Zambian data with windows of size 801 SNPs down sampled to 401 SNPs (A). For the Raleigh data, we used windows of size 401 SNPs (B). Each data point represents the H12 value calculated in an analysis window centered at the particular genomic position. Grey points indicate regions in the genome with recombination rates lower than 0.5 cM/Mb excluded from our analysis. Orange bard indicate common inversions present in at least in strains in each data set. There are multiple overlapping inversions on chromosome 3R. Red and blue points highlight the top 25 H12 peaks in the each respective scan relative to the median H12 value in the data. Red points indicate peaks that overlap both scans. The three highest peaks in the Raleigh scan correspond to known cases of soft sweeps at *Ace, CHKov1,* and *Cyp6g1*. We also observe peaks at *Ace* and *Cyp6g1* in the Zambian scan, but these do not rank as highly because the adaptive alleles at these loci are at lower frequencies at these sites.

To identify individual selective sweeps in the Zambian and Raleigh data, we found H12 peaks, where a peak is defined as a consecutive tract of H12 windows lying above the median H12 value in the data (Garud *et al*. 2015) (Methods). To focus on the most extreme selected events, we highlighted the top 25 H12 peaks in each scan in Figure 2. We excluded any peaks within 500Kb of the highest peaks to avoid identifying multiple peaks in the same region belonging to the same selective event (Methods). As previously observed in Figures 1c, 1d and S3, the H12 values at the peaks in the Zambian data are lower overall than in the Raleigh data. Reflecting this lower level of H12 in Zambian data, in Figure 2 the Zambian scan has a shorter y-axis than the Raleigh scan to have better resolution to identify individual selection events, but we show this data again in Figure S5 with the same y-axis as in Raleigh.

H12 peaks are distributed throughout the genome in both the Zambian and Raleigh populations, suggesting that no single region of the genome is driving the elevation of H12 values observed in the data. Three of the top 25 peaks in the Zambian scan overlap in location with three of the top 25 peaks in the Raleigh scan, where an overlap is defined as the intersection of the end coordinates of a peak (see Methods). This is greater than expected by chance (one sided binomial probability < 0.022, see Methods for calculations). One of the overlapping peaks coincides with the positive control *Cyp6g1* on chromosome 2R, and two of the overlapping peaks are on chromosome 3R. However, we find that these two peaks on chromosome 3R disappear when we rerun the scan in 801 SNP windows in Zambian data in Figure S4. Furthermore, we find an enrichment of peaks on chromosome 3R overlapping high frequency inversions in the scan of Zambian data in Figure 2 run with 801 SNP windows down sampled to 401 SNPs (see Table S4 for the enrichment calculations). This suggests that the two peaks in the Zambian scan overlapping peaks in the Raleigh scan on chromosome 3R may be either artifacts resulting from the presence of inversions or may be weakly selected events, and that only the peak at *Cyp6g1* is the only identified peak among the top 25 peaks present in both populations.

In addition, we identified six peaks on chromosome 3L in the Zambian population in contrast to just one peak in the Raleigh data, suggesting that there may be different selective pressures and different sources of adaptive mutations giving rise to the different peaks throughout the genome in the two populations.

We recovered H12 peaks at the positive controls *Cyp6g1* and *Ace* in both the Raleigh and Zambian data sets. While peaks at *Cyp6g1* and *Ace* rank 1^st^ and 3^rd^ in the Raleigh scan, they rank 14^th^ and 35^th^ in the Zambian scan. The lower rank of the peaks at *Ace* in the Zambian scan reflects the lower frequencies at which the adaptive mutations at *Ace* are present in the Zambian and Raleigh data respectively: approximately 16% in the Zambian and approximately 40% in the Raleigh populations. The second-ranking peak in the Raleigh data is at the positive control *CHKov1,* reiterating our previous result (Garud *et al*. 2015). However, this peak is absent in the Zambian scan, consistent with the previous observation that the adaptive mutation at *CHKov1* is present at high frequencies in out-of-Africa populations and rare in Africa (Aminetzach *et al*. 2005).

### Haplotype structure at the top H12 peaks in the Zambian and Raleigh scans

To gain intuition as to whether the top peaks in the H12 scans of the Raleigh and Zambian data sets resemble hard or soft sweeps, we visualized the haplotype frequency spectra in the analysis window with the highest H12 value at each of the top 25 peaks (Figures 3 and S4). The number of independent haplotypes present at high frequency in an analysis window can be associated with the ‘softness’ of a sweep. A substantial number of peaks in the H12 scan of the Zambian data show multiple haplotypes at high frequency, consistent with a signature of a soft sweep. In some cases, a few peaks look only borderline soft with one haplotype at high frequency followed by several haplotypes common to a few individuals in the population. In the H12 scan of the Raleigh data, all peaks have multiple haplotypes at high frequency resembling soft sweeps, as we previously observed (Garud *et al*. 2015). The sweeps at the top peaks in the Zambian data set reach a smaller partial frequency than in the Raleigh data set, again consistent with the muted elevation in linkage disequilibrium and haplotype homozygosity in the Zambian data set relative to the Raleigh data set.

**Figure 3:**
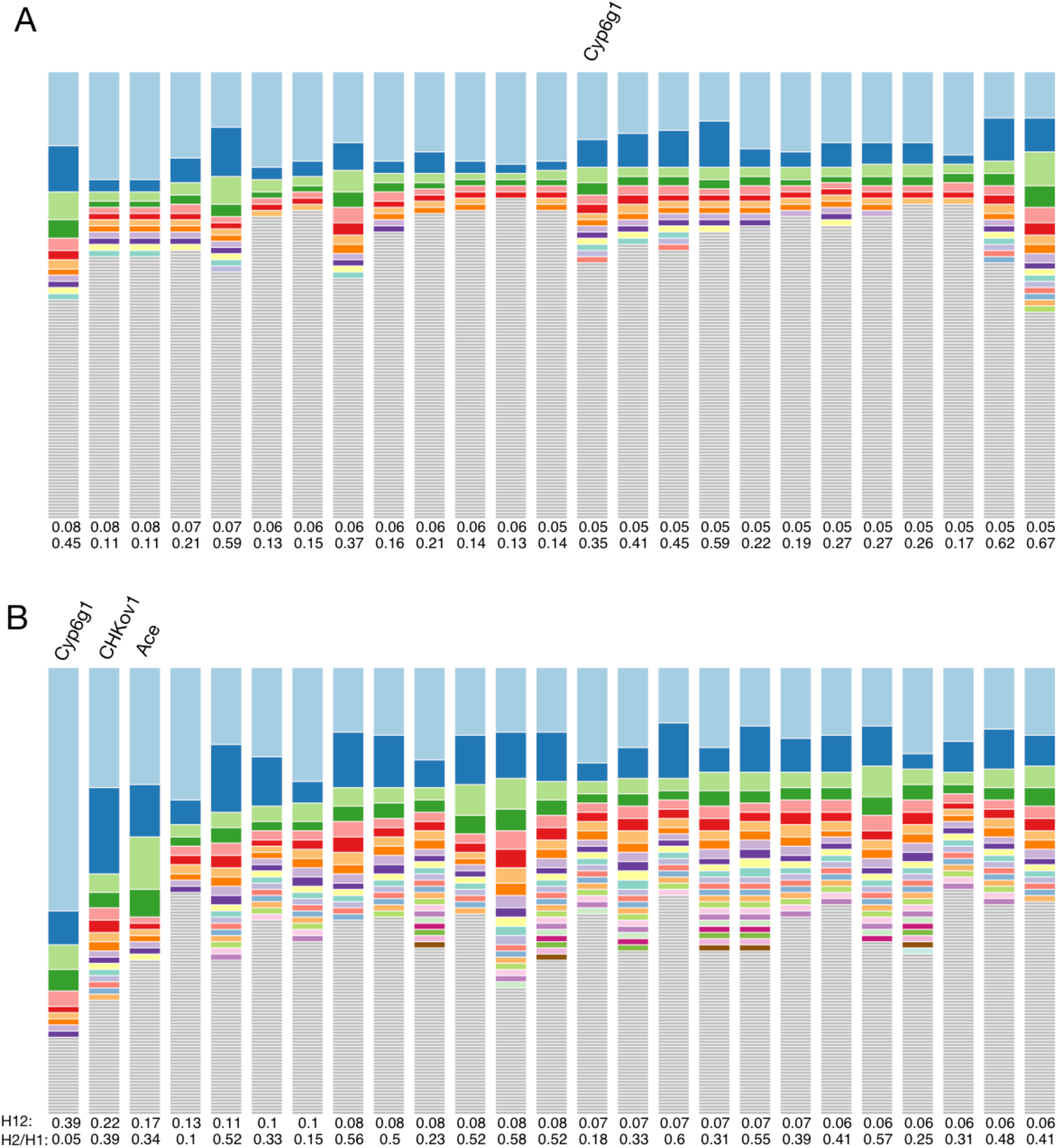
Haplotype frequency spectra for the 25 H12 peaks in Zambian and Raleigh data. Shown are haplotype frequency spectra for the top 25 peaks in the Zambian scan conducted in 801 SNP windows down sampled to 401 SNPs (A) and the Raleigh scan conducted in 401 SNP windows (B). For each peak, the frequency spectrum corresponding to the analysis window with the highest H12 value is plotted. The height of the first bar (light blue) in each frequency spectrum indicates the frequency of the most prevalent haplotype in the sample of 145 individuals, and heights of subsequent colored bars indicate the frequency of the second, third, and so on most frequent haplotypes in a sample. Grey bars indicate singletons. In Zambia, sweeps reach a smaller partial frequency than Raleigh. Most peaks in the Zambian data have multiple haplotypes present at high frequency indicative of soft sweeps, but there are also a few sweeps that look hard. In Raleigh all sweeps have multiple haplotypes at high frequency and look soft.

### Could demographic processes generate the bulk of the haplotype structure in Zambian data?

We tested whether the elevation in haplotype frequencies among the top 25 peaks in the Zambian data set could be generated by demographic processes such as admixture or backflow of individuals from North America to Zambia.

First, we compared genome-wide admixture levels in strains comprising the 3 most common haplotypes versus strains comprising all other haplotypes in the analysis window with the highest H12 value at each of the 25 peaks. We focused on the 3 most common haplotypes as these contribute to the bulk of the H12 values and we used admixture tracts inferred by Pool *et al*. (2012) for each Zambian strain for our analysis. We found that of the 25 peaks tested, only 1 peak, located on chromosome 3R, had significant differences in levels of admixture (P-value = 0.01743997, Wilcoxon two sided test), consistent with the expected number of false positives in 25 tests with a 0.05 significance level.

We also tested for any underlying population structure in the Zambian data by examining whether the same strains comprise the top 3 haplotypes more often than expected by chance across the top 25 peaks. We observed 15 strains comprising the top three haplotypes at 9 or more peaks, though not necessarily the same 9 peaks (P-value = 0.0017, see Methods for permutation analysis). This suggests that some of the long-range linkage disequilibrium observed in Figure 1 can be explained by some low level structure in the data, but that the majority of the haplotype homozygosity at the peaks are not driven by the same strains.

Additionally, we tested the possibility of backflow of flies from North America to Zambia. While African and European flies founded the North American population of flies (Duchen *et al*. 2013), the reverse scenario of migration from North America to Africa could have occurred (backflow). If such a migration event occurred, then common haplotypes in North America could be responsible for generating H12 peaks in the Zambian data. To test this, we compared diversity (π) in the less stringently filtered Raleigh data set in regions corresponding to the locations of top 25 peaks in the Zambian data versus all other regions in the Raleigh data. We found that the mean π in the two types of regions are similar (P-value = 0.5393864, see Methods).

To further test the backflow hypothesis, we investigated whether any of the top three common haplotypes among the 25 peaks in Zambian data are also common in Raleigh data. To identify shared common haplotypes, we aligned the start and end positions of the analysis windows with the highest H12 values in each of the 25 Zambian peaks with the less stringently filtered Raleigh data set and looked for matching haplotypes in both the Raleigh and Zambian data sets. We found only one peak with a common haplotype shared among 9 Zambian individuals and 8 Raleigh individuals. This peak is at the location of the *Cyp6g1* selective sweep, suggesting that either parallel adaptation or some migration event followed by adaptation in both Zambia and Raleigh has occurred at this locus.

Finally, we also tested the reverse scenario of whether H12 peaks identified in the Raleigh data are enriched in ‘ancestry disequilibrium hubs’ as identified by Pool 2015. An ancestry disequilibrium hub is characterized as a cluster of several African-African or European-European pairs of SNP with higher than expected levels of LD, suggesting that African-European SNP pairs are less compatible. We found that 2 of the top 25 peaks identified in the Raleigh scan overlap the ancestry disequilibrium hubs (P-value: 0.2820319, binomial, see methods).

Given the lack of evidence for backflow or admixture driving the elevation in haplotype homozygosity at the top peaks in the Zambian data, it is likely that the majority of H12 peaks observed in the Raleigh and Zambian data sets are driven by independent events. Thus, common soft sweeps and elevated LD are common to both populations. At the positive control, *Cyp6g1,* we saw that either parallel adaptation or migration followed by adaptation may have occurred. In the next section, we will see that haplotype sharing is common at all positive controls, *Ace, Cyp6g1* and *CHKov1*.

### Haplotype sharing in the Zambian and Raleigh populations at *Ace, Cyp6g1,* and *CHKov1*

We investigated whether common haplotypes are shared in both Zambia and Raleigh populations at all the positive controls, *Ace, Cyp6g1* and *CHKov1*. To identify shared haplotypes, we examined haplotype frequency spectra in 801 SNP windows at the sweep centers in a joint data set consisting of all Zambian and Raleigh strains from the less stringently filtered data sets (Methods). 801 SNP windows correspond to a slightly smaller analysis window size in terms of basepairs on average than in the Raleigh or Zambian data alone. In Figure 4, we show the haplotype frequency spectra at the positive controls in the joint data set and demarcate which population each haplotype is from.

**Figure 4:**
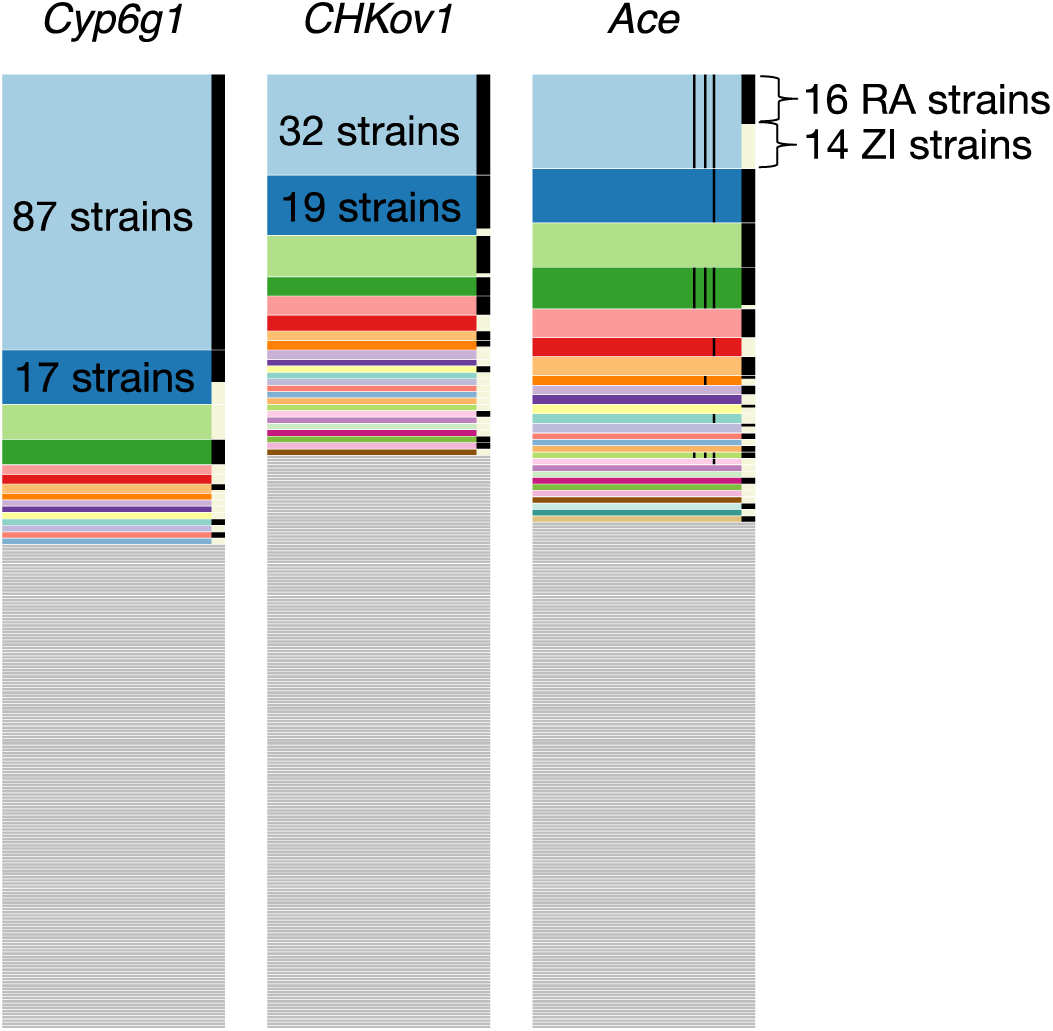
Haplotype frequency spectra at *Cyp6g1, CHKov1,* and *Ace*. Shown are haplotype frequency spectra at the three positive controls in a joint data set comprised of 300 Raleigh strains and Zambian strains in 801 SNP windows. On the right side of each of the frequency spectra are black and white bars indicating which strains are from Raleigh (RA) and Zambia (ZI), respectively. At all three positive controls common haplotypes are shared across the two populations. The thin black lines shown in the haplotype spectrum for *Ace* correspond to the presence of three adaptive mutations that confer resistance to pesticides at *Ace*. Three haplotypes have all three adaptive mutations. Raleigh and Zambian strains share two of these haplotypes.

At *Ace*, a common haplotype bearing three adaptive mutations previously discovered to confer resistance to pesticides in both Zambia and Raleigh is shared between 14 strains from Zambia and 39 strains from Raleigh. Another common haplotype in the Raleigh data comprised of 17 strains matches 1 Zambian strain and also bears the three adaptive mutations. The sharing of these haplotypes support the finding by Karasov *et al*. 2010 where it was observed that North American and Australian *D. melanogaster* strains share a haplotype bearing the three adaptive mutations, suggesting an out-of-Africa origin for this haplotype. Our finding provides a more complete adaptation history of *Ace*. All remaining high frequency haplotypes are comprised exclusively of either Raleigh or Zambian strains, many of which bear one or three adaptive mutations, suggesting that this soft sweep has arisen multiple times on multiple continents.

At the *CHKoV1* locus, we observe two common Raleigh haplotypes matching rare Zambian haplotypes: 17 Raleigh individuals and 2 Zambian individuals share one haplotype and 12 Raleigh individuals and 1 Zambian individual share another haplotype. Additionally, a third haplotype is shared between 1 Raleigh individual and 2 Zambian individuals. The sharing of common Raleigh haplotypes with rare Zambian haplotypes is consistent with the result in Aminetzach *et al*. 2005 that the adaptive allele at *CHKoV1* is rare in African populations and common in derived populations.

At *Cyp6g1,* we observe a common haplotype shared in 10 Raleigh individuals and 7 Zambian individuals (the exact frequencies in this analysis are slightly shifted from the earlier values in the previous section of 9 and 7 strains, respectively, because the analysis window is centered around the known selected locus rather than the analysis window with the highest H12 value). All other haplotypes are comprised exclusively of Raleigh or Zambian strains. This result suggests that this sweep has either arisen multiple times on different continents, and/or may share a common origin, similar to the sweep at *Ace*.

### Bayesian analysis of the softness of the peaks

To assess whether the top peaks in the two scans can be more easily generated under simulated hard and soft sweeps, we measured H2/H1 (Garud *et al*. 2015) in the windows with the highest H12 values at each of the top peaks. H2/H1 is a ratio of haplotype homozygosity calculated without (H2) and with (H1) the most frequent haplotype in the sample. This statistic, used in conjunction with H12, can distinguish hard and soft sweeps as long as the sweeps are not too soft (Garud *et al*. 2015). A sweep that is too soft to be detected by H12 and differentiated by H2/H1 as hard or soft has such a high number of unique haplotypes bearing an adaptive mutation that the sweep cannot be distinguished easily from neutrality (Garud *et al*. 2015).

We performed a Bayes factor analysis to assess the likelihood of a hard versus soft sweep generating a particular pair of H12 and H2/H1 values for each of the top peaks. Bayes factors are calculated as follows: BF = P(H12_obs_, H2_obs_ /H1_obs_ | Soft Sweep)/P(H12_obs_, H2_obs_ /H1_obs_ | Hard Sweep). We approximated BFs using an approximate Bayesian computation (ABC) approach similar to our previous analysis (Garud *et al*. 2015) where the parameters selection coefficient (s), partial frequency of the adaptive allele after selection has ceased (*PF*), and age (*T_E_*) are drawn from uniform prior distributions: *s* ~ U[0,1], *PF* ~ U[0,1], and *T_E_* ~ U[0,0.001]*×N_e_*. Similar to our previous analysis, we simulated hard sweeps with an adaptive mutation rate *θ*_A_ = 0.01 and soft sweeps with *θ*_A_ = 10. To assess the softness of the top 25 peaks in the Zambian data, we performed simulations with a constant *N_e_*=4.8*10^6^ demographic model (matching the inferred *N_e_* for the Zambian population, Methods) and measured H12 and H2/H1 in windows of 801 SNPs down sampled to 401 SNPs. To assess the softness of the top 25 peaks in the Raleigh data, we performed simulations with a constant *N_e_*=2.6×10^6^ demographic model (matching the inferred *N_e_* for the Raleigh population, Methods) and measured H12 and H2/H1 in windows of 401 SNPs.

In Figure 5 we present panels of Bayes factors measuring the relative support of hard versus soft sweeps for a grid of different values of H12 and H2/H1. A substantial number of the top 25 peaks in the Zambian data have H12 and H2/H1 values that are exceptionally well supported by soft sweeps. A few peaks have H12 and H2/H1 values that are borderline supported by soft sweeps. However, the values of H12 at all peaks in the Zambian data are low, and as we have previously shown (Garud *et al*. 2015), it is difficult to differentiate hard and soft sweeps at peaks with low H12. This muted pattern in Zambia is in contrast to the top 25 peaks in the Raleigh data which all have high H12 and H2/H1 values that are well supported by soft sweeps, we have seen previously (Garud *et al*. 2015). Overall the substantial number of peaks with H12 and H2/H1 values consistent with soft sweeps in both populations suggest that soft sweeps are a generic feature of multiple populations of *D. melanogaster*.

**Figure 5:**
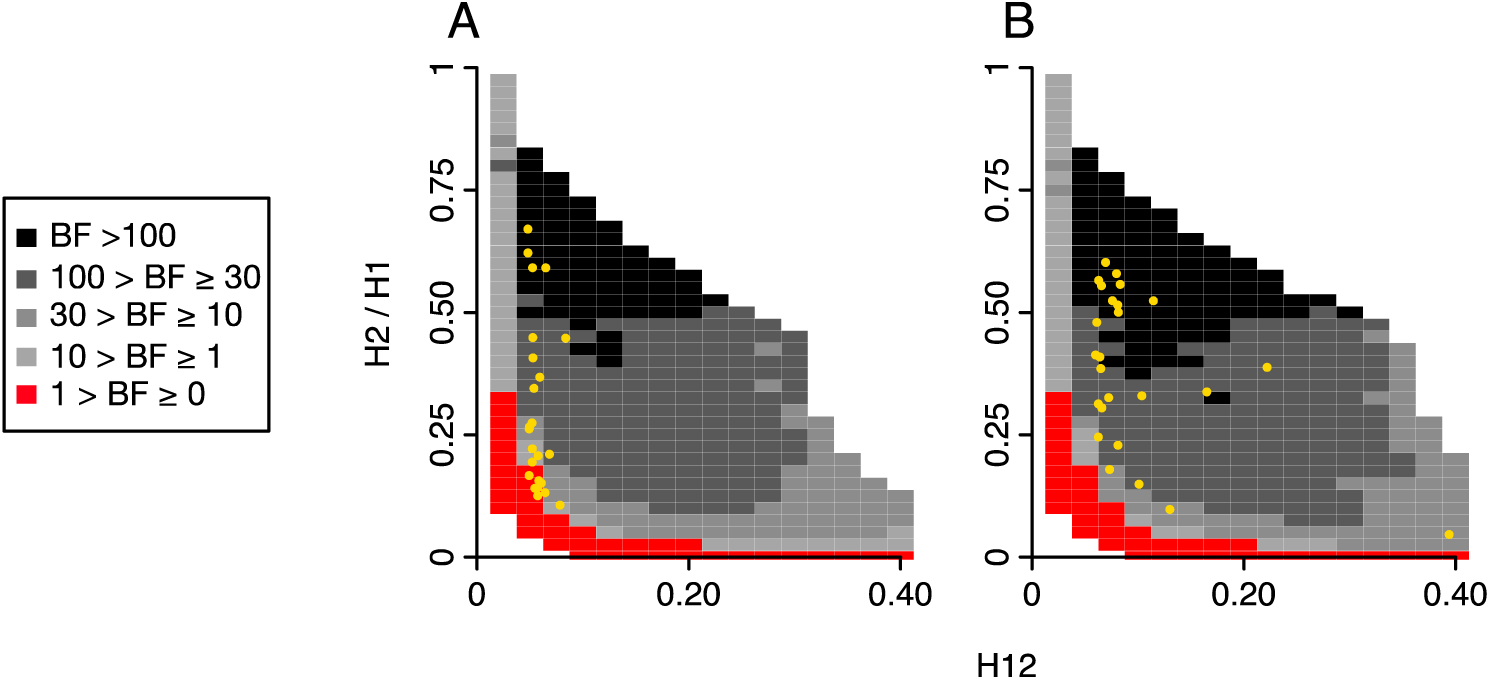
Range of H12 and H2/H1 values expected under hard and soft sweeps for Zambian and Raleigh data. Bayes factors (BFs) were calculated for a grid of H12 and H2/H1 values to demonstrate the range of H12 and H2/H1 values expected under hard versus soft sweeps for the top 25 peaks in the Zambian (A) and Raleigh (B) scans. BFs were calculated by taking the ratio of the number of soft sweep versus hard sweep simulations that were within a Euclidean distance of 10% of a given pair of H12 and H2/H1 values. Red portions of the grid represent H12 and H2/H1 values that are more easily generated by hard sweeps, while grey portions represent regions of space more easily generated under soft sweeps. Each panel presents the results from 10^6^ hard sweep simulations generated with *θ*_A_ = 0.01 and 10^6^ soft sweep simulations generated with *θ*_A_ = 10. For any colored square, at least 1000 hard and soft sweep simulations match the corresponding pair of H12 and H2/H1 values of the grid. White portions of the grid represent cases where fewer than 1000 simulations match the pair of H12 and H2/H1 values. Simulations were performed in samples of 145 individuals with a recombination rate (*ϱ*) of 0.5 cM/bp under a constant *N_e_*=4.8*10^^^6 demographic model for comparison with the Zambian data (A) and a constant *N_e_=*10^6^ demographic model for comparison with the Raleigh data. Overlaid on each plot in yellow are the observed H12 and H2/H1 values for the top 25 peaks for each scan. The H12 and H2/H1 values measured at the top 25 peaks in Zambian data can be generated by both hard and soft sweeps, whereas the H12 and H2/H1 values measured in Raleigh can be more easily generated by soft sweeps than hard sweeps in all scenarios.

## DISCUSSION

We previously showed that patterns of elevated LD and haplotype homozygosity consistent with soft sweeps are common in *D. melanogaster* strains from Raleigh (Garud *et al*. 2015). Here we compare these patterns with that of a second population from Zambia to test the extent to which this signal in the Raleigh population may be driven by peculiarities of the data such as the extensive admixture of the Raleigh population or the inbreeding performed to generate the sequenced lines. Unlike the Raleigh population, the Zambian population has undergone relatively little admixture, and instead of inbreeding lines, the data set was generated by sequencing haploid embryos.

We find that both LD and haplotype homozygosity is elevated at all distances in both Zambian and Raleigh data relative to expectations in several neutral demographic models that match *S* and *π* estimates in the data. This elevation remains despite controlling for inversions, admixture, regions of low recombination, relatedness between individuals, missing data, sample size, and allele frequency. Furthermore, we find that signatures of soft sweeps are common in both the Zambian and Raleigh data sets. We tested for the possibility of admixture or backflow generating the haplotype structure observed in Zambian data, but found that the signatures observed in this population are independent of the Raleigh population, with the exception of specific cases at the positive controls, *Ace, Cyp6g1,* and *CHKov1*. The extent of the elevation in LD and haplotype homozygosity and the number of soft sweeps in both populations is surprising and simultaneously confirms that these results are generic to multiple populations of *D. melanogaster*.

Yet, there are some differences in the patterns observed in the Zambian and Raleigh data set. For example, the elevation in LD and haplotype homozygosity is muted in the Zambian population relative to the Raleigh population and all the selective sweeps in Zambia have lower partial frequencies and H12 values than in Raleigh. In addition, while most of the top 25 peaks’ H12 and H2/H1 values in the Zambian data are overwhelmingly supported by soft sweeps, a few of the top peaks are borderline soft with one haplotype at high frequency and several lower frequency haplotypes present in a few individuals. It is difficult to differentiate whether these sweeps are truly hard or soft because H2/H1 has poor ability to differentiate the softness of a sweep with a low H12 value (Garud *et al*. 2015). The exaggerated signals in the Raleigh data set could in principle be driven by the inbreeding and admixture experienced by the Raleigh population, but it is clear that these two forces alone cannot explain all the elevation in haplotype homozygosity and LD given that these signatures are generic to both populations.

Instead, another possibility why the signals in the Raleigh population are more pronounced could be that Raleigh has experienced stronger and more recent selection pressures, especially due to anthropogenic pressures such as those arising from the presence of pesticides. We can extrapolate from the positive controls at *Ace, Cyp6g1,* and *CHKov1,* cases of recent and strong positive selection arising from the presence of pesticides and viruses, to learn more about the strength and timing of the selection pressures in Zambia versus Raleigh. Examining the frequencies of the adaptive mutations at the positive control *Ace* shows that the adaptive mutations have reached frequencies of ~40% in Raleigh and only ~16% in Zambia. The peaks at *Cyp6g1* and *Ace* rank 14^th^ and 35^th^ in the H12 scan in Zambia, in contrast to 1^st^ and 3^rd^ in the Raleigh scan likely because the adaptive mutations have reached lower frequencies in Zambia. We do not even observe at peak at *CHKov1* in Zambia, consistent with the finding in Aminetzach *et al*. 2005 that the adaptive allele is rare in Africa and common only in derived populations at this locus.

The haplotype patterns at *Ace, Cyp6g1,* and *CHKov1* in both Zambian and Raleigh data show that these soft sweeps have arisen independently on different continents as well as from single origins spread by migration. At all three loci, multiple haplotypes at high frequencies are unique to both the Zambian and Raleigh populations, showing that independent soft sweeps are common in different locations around the world. Additionally, at all three loci, we observed that a common haplotype is shared by both Zambia and Raleigh strains, suggesting that migration also plays a role. At *Ace,* the shared common haplotype bears three adaptive mutations in contrast to haplotypes bearing the single adaptive mutation present at high frequencies comprised uniquely of Zambia or Raleigh strains. This suggests that this haplotype arose once in Africa and spread world wide, confirming the results of Karasov *et al*. 2010. At *Cyp6g1* we find for the first time that a common haplotype is shared in both Zambia and Raleigh. At *CHKov1* we find that common haplotypes in Raleigh are shared with rare haplotypes in Zambia, consistent with the result in Aminetzach *et al*. 2005 showing that the adaptive allele originated in Africa and rose to high frequencies in derived population.

Intriguingly, we notice that virtually all sweeps in Raleigh and Zambia reach an intermediate frequency and none go to completion. it is possible that *D. melanogaster* experiences a high genome-wide rate of partial selective sweeps, contributing to the elevation in LD. Current McDonald-Kreitman estimates of the rate of adaptation (Andolfatto 2005; Bierne and Eyre-Walker 2004; Eyre-Walker and Keightley 2009; Fay *et al*. 2002; Kousathanas and Keightley 2013; Schneider *et al*. 2011; Sella *et al*. 2009; Shapiro *et al*. 2007; Smith and Eyre-Walker 2002) may be lower than the true adaptive rate if partial sweeps are common and often fail to go to fixation in the long term because McDonald-Kreitman estimates measure fixations of polymorphisms over very long time scales. This will need to be tested further with forward simulations of realistic demographic and selection scenarios.

Although we tested several neutral demographic scenarios fit to the Zambia and Raleigh data, we could not match the elevation in LD and haplotype homozygosity observed in the data. It is possible that there is a demographic scenario yet to be identified that can generate the observed levels of LD and haplotype homozygosity. One such extreme scenario that could generate the patterns observed is an admixture model where two parental populations undergo sharp bottlenecks resulting in virtually isogenic populations followed by an expansion before mixing. Alternatively, there could be demes within each population generating structure in the data that we are unable to model. Going forward, simulations of more realistic demographic scenarios potentially also incorporating selection are needed to fully understand the underlying forces in these populations.

Although background selection and gene conversion have been shown to act pervasively throughout the *D. melanogaster* genome (Comeron 2014; Comeron *et al*. 2012), it is unlikely that these two forces significantly influence the patterns of LD and haplotype homozygosity: while background selection can depress levels of polymorphisms, on its own it cannot elevate haplotype to high frequencies (Enard *et al*. 2014; Fagny *et al*. 2014). Gene conversion has been shown to act on short scales of 50 to 2000 bps at most (Hilliker *et al*. 1994; Jeffreys and May 2004) and should thus only impact LD on short scales and not long-range LD. In fact, it has been noted that gene conversion should decrease LD on short scales, thereby making haplotype homozygosity tests that do not take into account gene conversion conservative (Andolfatto and Nordborg 1998).

Long range LD has been observed in many other organisms in addition to *D. melanogaster* including cattle, wheat, maize, sheep, and humans (Farnir *et al*. 2000; Maccaferri *et al*. 2005; Miller *et al*. 2011; Palaisa *et al*. 2004; Pritchard and Przeworski 2001; Rybicki *et al*. 2002). It has been shown that there are varied sources of LD in these different organisms including strong selection, admixture events, and population substructure. Thus the evolutionary forces explored in this study of LD and haplotype homozygosity in *D. melanogaster* are not unique. It is becoming increasingly clear that in order to better understand the full spectrum of forces shaping natural variation in multiple populations, selective and demographic forces need to be inferred jointly.

## METHODS

### Quality filtering of the Raleigh and Zambian data

The Raleigh strains used for the analysis were generated by Mackay *et al*. 2012 and belong to the Drosophila Genetic Reference Panel. The DGRP data set consists of fully sequenced genomes of 197 inbred *D. melanogaster* lines collected from Raleigh, North Carolina. The Zambian strains belong to the Drosophila Genome Nexus (Lack *et al*. 2015) and consist of 205 fully sequenced strains.

We masked regions in the data inferred to have high levels of identify by descent (IBD), admixture, and inversions because these are all factors that can contribute to high LD. Scripts for masking were provided by John Pool (www.johnpool.net) along with regions inferred to have high IBD and admixture (Lack *et al*. 2015). Only African strains were masked for admixture between African and European strains. North American strains have been shown to have a high level of genome-wide admixture (Duchen *et al*. 2013), and so admixture in these strains were controlled for by performing neutral simulations under the admixture model by Duchen *et al*. 2013. All heterozygous sites and inferred tracts of residual heterozygosity (Lack *et al*. 2015) were masked as well.

The extensive masking resulted in large tracts of ‘Ns’ in the data with an average length of 4MB. In some cases, entire chromosomes were masked. However, strains with high levels of masking on one chromosome did not necessarily have high levels of masking on other chromosomes. To account for this heterogeneity in the strains with masking on each chromosome, we down sampled to 100 strains with the least number of Ns per chromosome. Thus, each chromosome consisted of different sets of 100 strains. The strain IDs used for each chromosome are given in Table S3.

To have the best resolution and power to detect individual selective sweeps with H12, we applied less stringent filters to the Raleigh and Zambian data sets and did not mask regions of IBD, admixture, and inversions. Instead, we excluded 8 Zambian and 27 Raleigh strains with genome-wide IBD levels with at least one other strain > 20% as calculated by Lack *et al*. 2015.

The excluded strain IDs are as follows: ZI397, ZI530, ZI269, ZI240, ZI218, ZI207, ZI523, ZI86RAL-385, RAL-358, RAL-712, RAL-399, RAL-879, RAL-355, RAL-810, RAL-350, RAL-832, RAL-882, RAL-306, RAL-799, RAL-801, RAL-859, RAL-907, RAL-790, RAL-748, RAL-336, RAL-850, RAL-365, RAL-786, RAL-730, RAL-861, RAL-59, RAL-646, RAL-812, and RAL-787. In this version of the data set, we masked all heterozygous sites and inferred tracts of residual heterozygosity as in the previous data set.

H12 was applied to the entire 178 Raleigh strains and 188 Zambian strains. To reduce any remaining heterogeneity in sample size due to missing data, we down sampled to the 145 strains with the least amount of missing data in each analysis window for inclusion in the H12 calculation.

### Demographic inference of the Zambian population

We fit four demographic models to the Zambian data using the program DaDi (Gutenkunst *et al*. 2009). DaDi calculates a log-likelihood of the fit of a model based on an observed site frequency spectrum (SFS). We estimated the SFS for presumably neutral SNPs in ZI data using segregating sites in short introns (Parsch *et al*. 2010). Specifically, we used every site in a short intron of length less than 86 bps, with 16 bps removed from the intron start and 6 bps removed from the intron end (Lawrie *et al*. 2013). We masked all IBD regions, admixture, inversions, and heterozygosity tracts in the Zambian data set. We did not exclude any strains, but rather, projected the SFS down to 100 samples at each polymorphic site. This procedure resulted in 73,197 SNPs in the Zambian data out of a total of 738,024 bps.

The models we fit to the Zambian short intron data include a bottleneck model, a growth model, a bottlegrowth model (a three epoch model with a bottleneck followed by a growth), and a constant *N_e_* model. For the bottleneck and bottlegrowth models, we inferred four free parameters: the bottleneck size (*N_B_*), the bottleneck time (*T_B_*), final population size (*N_F_*), and the final population time (*T_F_*) (Table S1). For the growth model, we inferred two free parameters: final population size (*N_F_*) and duration of the growth (*T*). Finally, we also inferred a constant *N_e_* = 4,789,108 (≈4.8×10^6^) for the constant population size model using a Watterson’s theta estimate of the Zambian data and assuming a mutation rate of 10^−9^/bp/generation. Similarly, we inferred a constant *N_e_* = 2,603,678 (≈2.6×10^6^) fit to Watterson’s theta estimate for the short intron data of Raleigh, similar to the constant *N_e_* model inferred by Garud *et al*. 2015.

### Simulations of neutral demographic scenarios and selection scenarios

Population samples under neutrality were simulated with the coalescent simulator MSMS (Ewing and Hermisson 2010). We simulated samples of size 100 to match the sample depth of the Raleigh and Zambia data and assumed a neutral mutation rate of 10^−9^ events/bp/gen (Keightley *et al*. 2009) and a recombination rate of 10^−6^ cM/bp.

We simulated population samples experiencing hard and soft selective sweeps with MSMS in sample sizes of 145 to match the sample depth of the data used for the H12 scans. We simulated hard sweep and soft sweeps arising from *de novo* mutations by specifying the population parameter *θ*_A_ = 4*N_e_u*_A_ at the adaptive site to be 0.01 and 10, respectively, matching the parameter estimates used in Garud *et al*. 2015. The adaptive site was always placed in the center of the locus. We performed 10^6^ hard and soft sweep simulations in constant *N_e_*=4.8*10^6^ and 2.6*10^6^ scenarios matching the expected constant *N_e_* population sizes for the Zambian and Raleigh populations, respectively. We assumed a neutral mutation rate of 10^−9^ events/bp/gen (Keightley *et al*. 2009) and a recombination rate of 5*10^−7^ cM/bp. We also assumed codominance, whereby a homozygous individual bearing two copies of the advantageous allele has twice the fitness advantage of a heterozygote. The selection coefficient (*s*), partial frequency of the adaptive allele after selection ceases (*PF*), and age (*T_E_*) were drawn from uniform prior distributions as follows: *s* ~ U[0,1], *PF* ~ U[0,1], and *T_E_* ~ U[0,0.001]×*N_e_*.

### Linkage disequilibrium estimates

We measured linkage disequilibrium (LD) in DGRP data and in neutral and selection simulations in samples of size 100. LD was measured using the *r*^2^ statistic in sliding windows of 1000 SNPs iterated by 50 SNPs. All 1000 SNP windows were greater than 10Kb in length. LD was measured between the first SNP in the window and the rest of the SNPs in the window with a distance between the SNPs less than or equal to 10Kb. If any SNP had missing data, the individuals with the missing data were excluded from the LD calculation as in Jakobsson *et al*. 2008. At least 4 individuals without missing data at both SNPs were required to compute LD, otherwise the SNP pair was discarded. If at least one SNP in a SNP pair resided in a region with recombination < 10^−6^ cM/bp, the SNP pair was discarded. LD plots were smoothed by averaging LD values binned in non-overlapping 20 bp windows until a distance of 300 bps. After that, LD values were averaged in bins of 150 bp non-overlapping windows. When controlling for minor allele frequency (MAF), both SNPs in each LD calculation were required to have a MAF in the interval desired.

### Genomic scan for selective sweeps in Zambian and Raleigh data using H12

We scanned the Zambian and Raleigh genomes with H12 for selective sweeps. For the Zambian data we used analysis windows of 801 SNPs randomly downsampled to 401 SNPs (Figure 2a) as well as 801 SNPs (Figure S3). For the Raleigh data, we used windows of 401 SNPs. The centers of the windows were all separated by intervals of 50 SNPs. Haplotypes were grouped together only if they matched at all sites. Any haplotype with missing data matching multiple haplotypes at all sites except the sites with missing data were randomly grouped with one other matching haplotype.

We identified H12 peaks in the data by grouping together consecutive tracts of H12 analysis windows lying above the median H12 value in the data. We chose the highest H12 value among all windows in the peak to represent the H12 value of the entire peak. We defined the edge coordinates of each peak as the smallest and largest coordinates of all SNPs comprising a peak across all windows. After identifying all peaks, we iterated through the peaks in reverse order starting with the highest H12 value and excluded all peaks within 500Kb from the center of each peak until all peaks had been accounted for. This avoided the potential problem of identifying multiple peaks in the same genomic region. Finally, we excluded peaks with edge coordinates overlapping any region identified by Comeron *et al*. 2012 with rho < 0.5 cM/Mb.

To identify overlapping peaks in the Zambian and Raleigh scans, we found the intersection of coordinate pairs for the peaks identified in the Zambian and Raleigh data sets. The overlapping peaks are highlighted in red in Figures 2 and S3.

To calculate the probability of 3 or greater overlapping peaks in the two scans, we calculated the fraction of each genome covered by H12 peaks. In Raleigh 1.51% of the genome is covered by H12 peaks (corresponding to 4,252,692 bps) and in Zambia 4.38% of the genome is covered by H12 peaks (corresponding to 1,461,431bps). The probabilities of observing 3 out of 25 peaks in Raleigh overlapping 3 out of 25 peaks in Zambia can be computed with a right-tailed binomial probability, where the probability of an overlap in Zambian or Raleigh data corresponds to either 1.51% or 4.38%, respectively. Thus the probability of 3 or more overlaps in the Raleigh data is 0.0224 and the probability of 3 or more overlaps in the Zambian data is 0.0005.

### Test for structure and backflow generating H12 peaks in Zambian data

We tested for an enrichment of strains among the top three haplotypes across all the peaks in the Zambian H12 scan. To do so, we shuffled the labels of strains comprising all the haplotypes at each peak and counted the number of strains appearing among the three most common haplotypes in *n* or more peaks, where *n* ranged from 0 to 25 peaks. We repeated this procedure 10,000 times. Then, comparing against the distribution of the number of repeated strains among the top 3 haplotypes for *n* or more peaks, we calculated an empirical P-value for observing 15 strains appearing among 9 or more peaks’ top haplotypes. We did the same for all other *n*s and found that for lower *ns* we did not get significant P-values.

We tested for the possibility of backflow of strains from North America to Zambia generating the H12 peaks in the Zambian data set. First we tested for a dip in diversity in the Raleigh data at positions corresponding to the top 25 peaks in the Zambian data set. We computed π in 10Kb non-overlapping windows on all autosomal arms in the Raleigh data. We then calculated the mean π in any window overlapping the start and end coordinates of each of the top 25 peaks identified in the Zambian data set and compared this mean with the distribution of π calculated in all remaining windows in the Raleigh data set.

### Test for enrichment of peaks in ancestry disequilibrium hubs

To test whether the top 25 peaks in the Raleigh data set are enriched in ancestry disequilibrium hubs identified by Pool 2015, we first identified that the end coordinates of the first and last windows of two peaks overlap the coordinates of the hubs listed in Table S6 of Pool 2015. We then calculated the fraction of the autosomes that the hubs occupy (8.1%) and then calculated a binomial probability of observing 2 out of 25 peaks overlapping any hub at random.

## ACKNOWLEDGEMENTS

We gratefully thank Philipp Messer, David Enard, Noah Rosenberg, Shaila Musharoff, and Alan Bergland for their invaluable help. This work was supported by the Stanford Center for Evolution and Human Genomics fellowship to NRG and NIH grants R01 GM100366, R01 GM097415, R01 GM089926 to DAP.

## SUPPLEMENT

**Figure S1:**
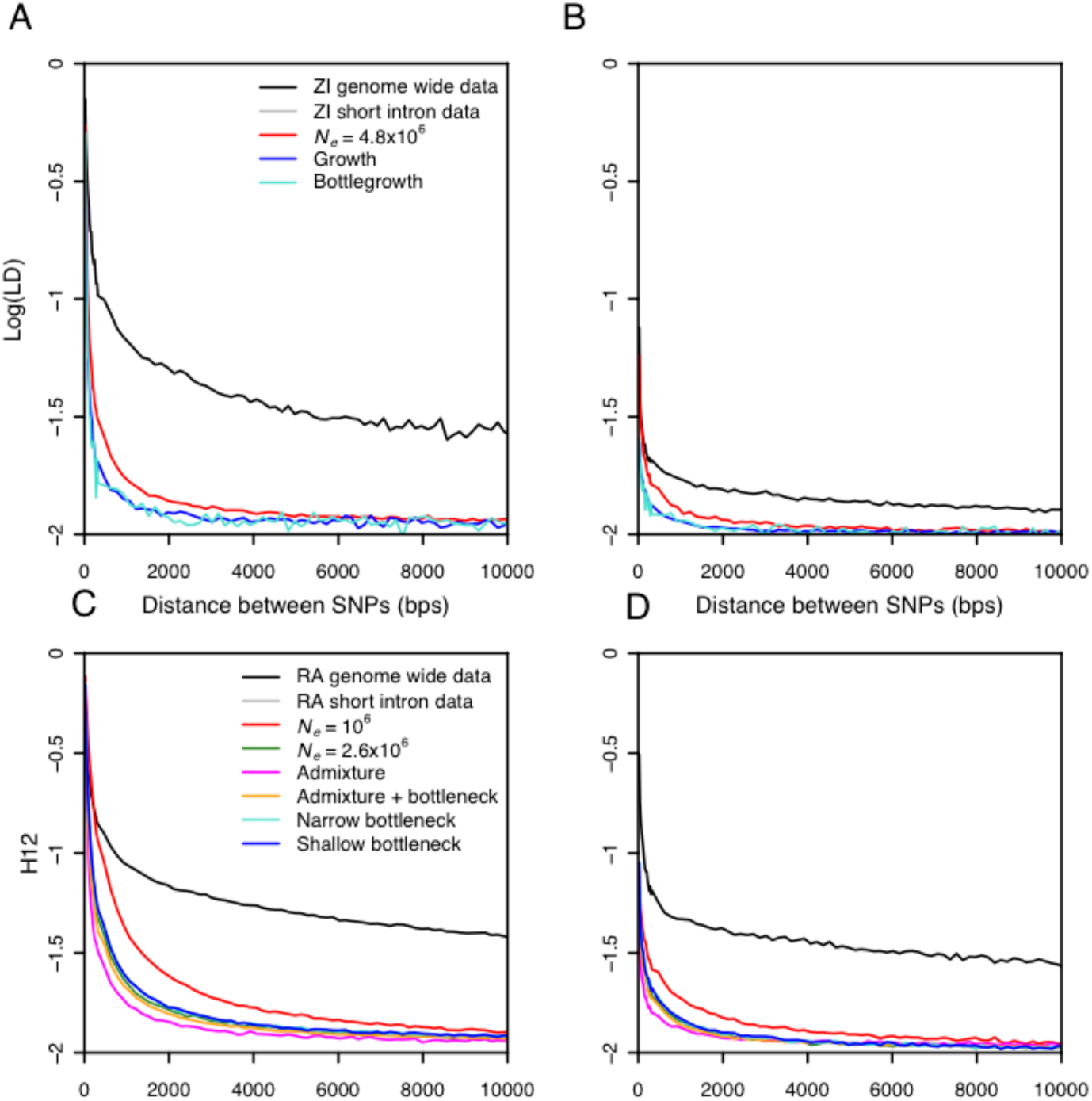
Elevation in LD values in Raleigh and Zambian data relative to neutral demographic models. In these panels, minor allele frequencies for SNPs were conditioned on values between 0.3 and 0.5, and 0.0 and 0.05. (A) Zambian (ZI) data and demographic models with MAFs between 0.3 and 0.5. (B) Zambian data and demographic models with MAFs between 0.0 and 0.05. (C) Raleigh (RA) data and demographic models with MAFs between 0.3 and 0.5. (D) Raleigh data and demographic models with MAFs between 0.0 and 0.05.

**Figure S2:**
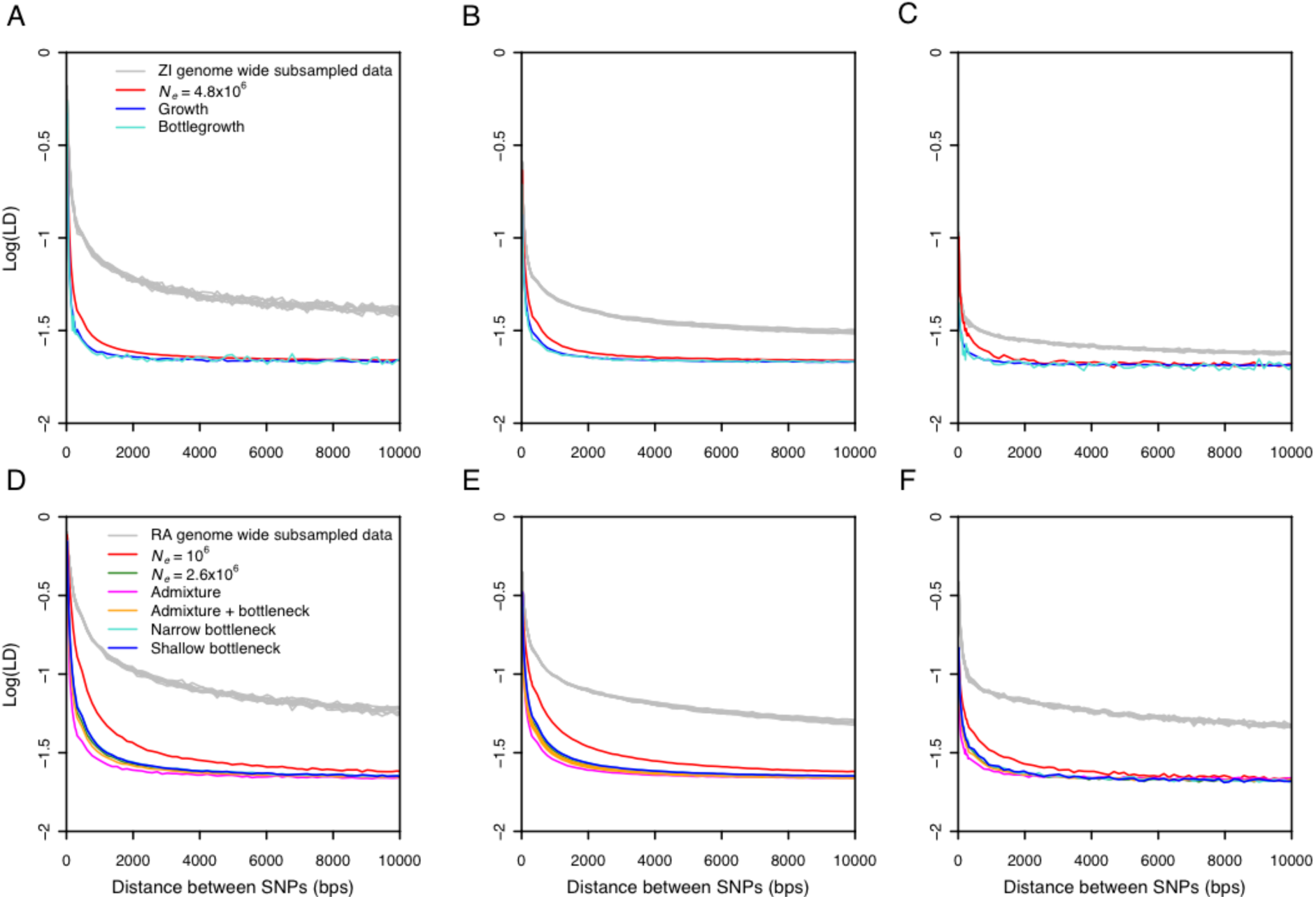
Elevation in LD values in Zambian and Raleigh data after down sampling to 50 strains. Zambian and Raleigh strains were down sampled to 50 strains 10 times and the resulting LD distributions were plotted (grey). In contrast to expectations under any neutral demographic model tested with a sample size of 50, all samples of 50 strains have elevated long range LD. This indicates that the elevation in LD is not driven by any sub-population. The LD calculations are conditioned on minor allele frequency (MAF) classes as follows: (A) Zambian (ZI) data and demographic models with MAFs between 0.3 and 0.5. (B) Zambian data and demographic models with MAFs between 0.05 and 0.5. (C) Zambian data and demographic models with MAFs between 0.0 and 0.05. (D) Raleigh (RA) data and demographic models with MAFs between 0.3 and 0.5. (E) Raleigh data and demographic models with MAFs between 0.05 and 0.5. (F) Raleigh data and demographic models with MAFs between 0.0 and 0.05.

**Figure S3:**
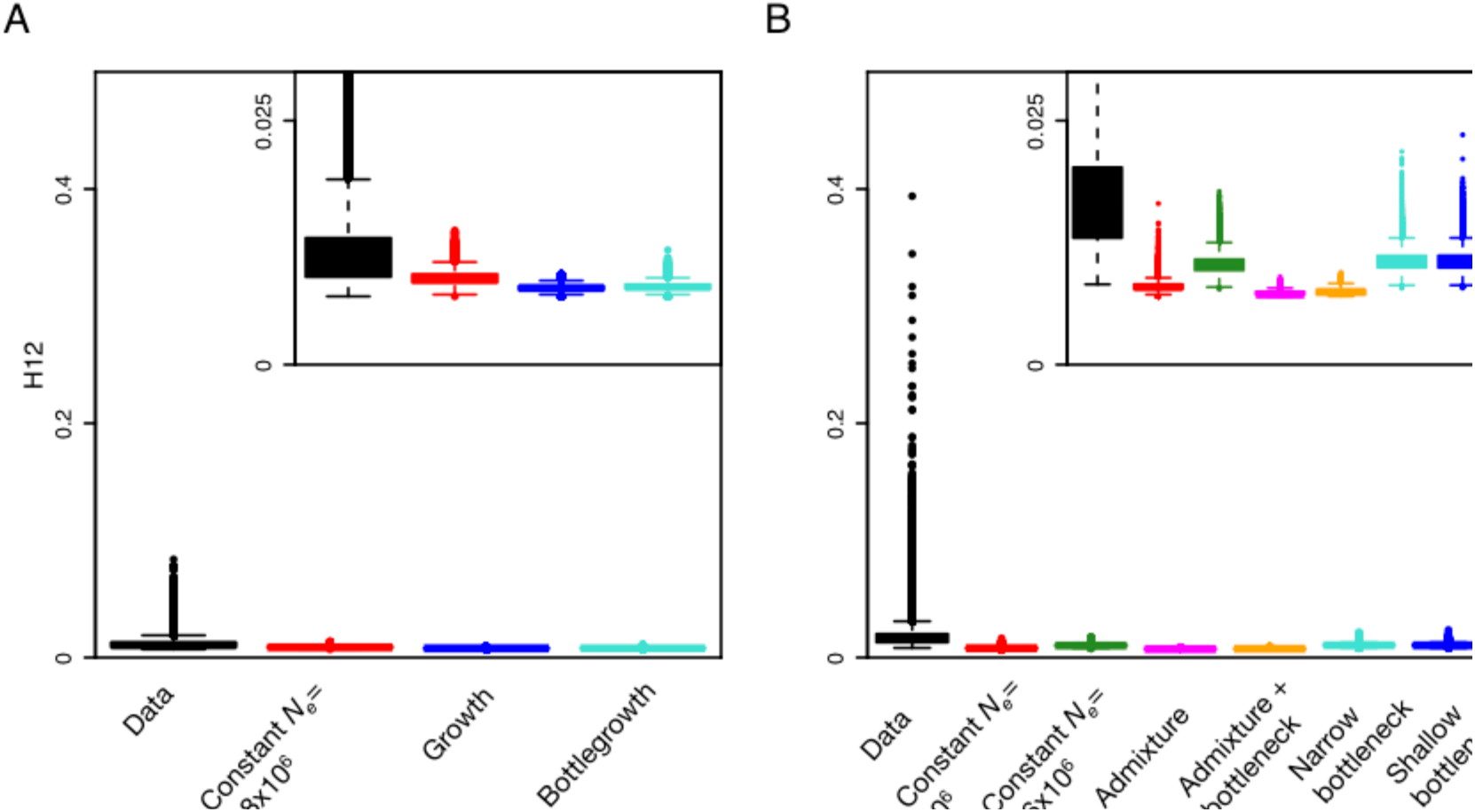
Elevation in H12 values in Zambian and Raleigh data with less stringent filters. Genome-wide H12 values measured in 801 SNP windows in Zambian data (A) and 401 SNP windows in Raleigh data (B) are elevated as compared to expectations under any neutral demographic model tested. In this data set, we did not mask inversions, IBD tracts, or admixture tracts. Instead we excluded individuals with > 20% of the genome sharing IBD tracts of 0.5 similarity or greater with any other another individual in the sample. H12 values were measured in sample sizes of 145 and in genomic regions with *ϱ ≥* 0.5 cM/Mb. H12 values were measured in neutral demographic simulations of 145 individuals generated with *ϱ* = 0.5 cM/Mb. Plotted are the result of approximately 1.5×10^5^ simulations under each neutral demographic model, representing ten times the number of analysis windows in the data.

**Figure S4:**
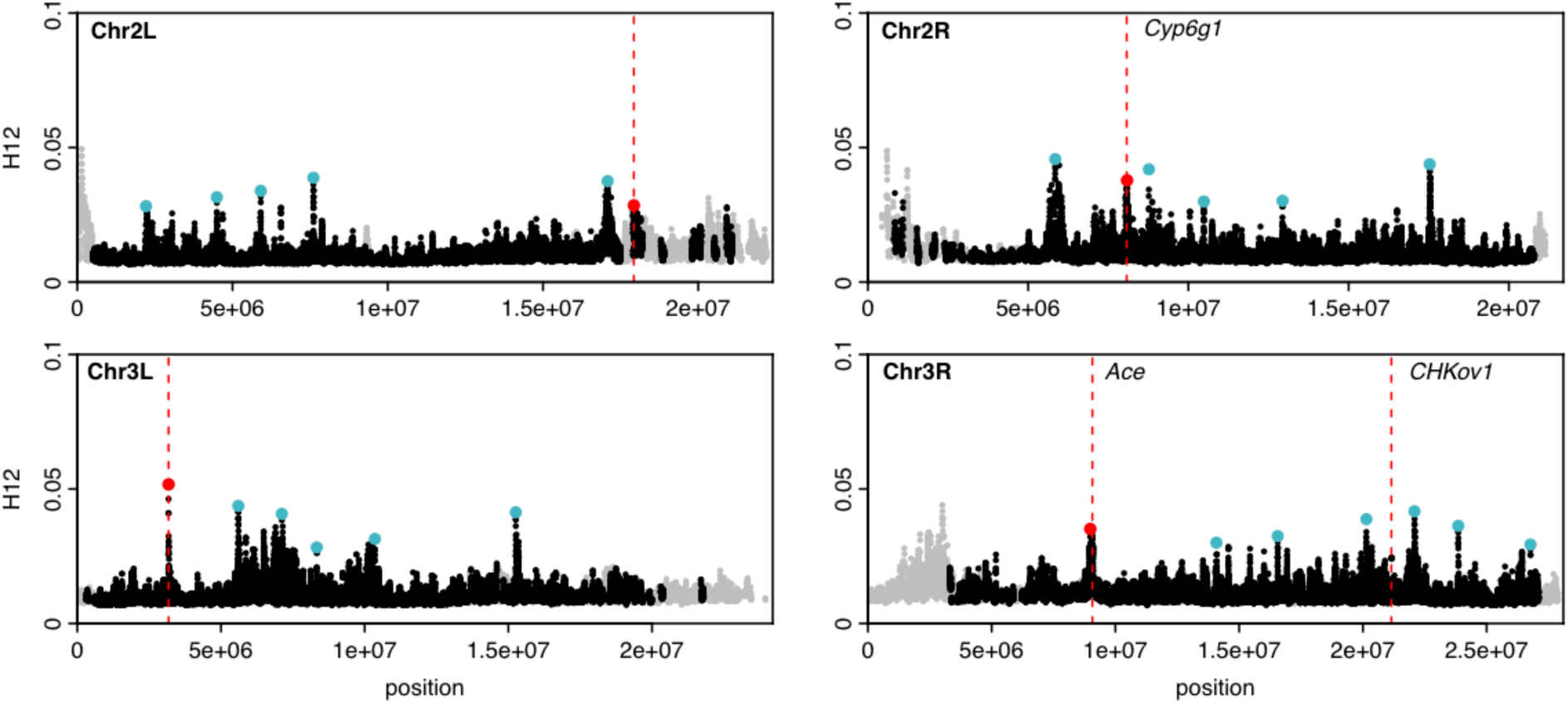
H12 scan in Zambian data in 801 SNP analysis windows. Plotted is an H12 scan conducted in Zambian data in 801 SNP windows. Grey points indicate regions in the genome with recombination rates < 0.5 cM/Mb excluded from the analysis. Red and blue points highlight the top 25 H12 peaks in the each respective scan relative to the median H12 value in the data. Red points indicate peaks that overlap the peaks in the H12 scan of Raleigh data.

**Figure S5:**
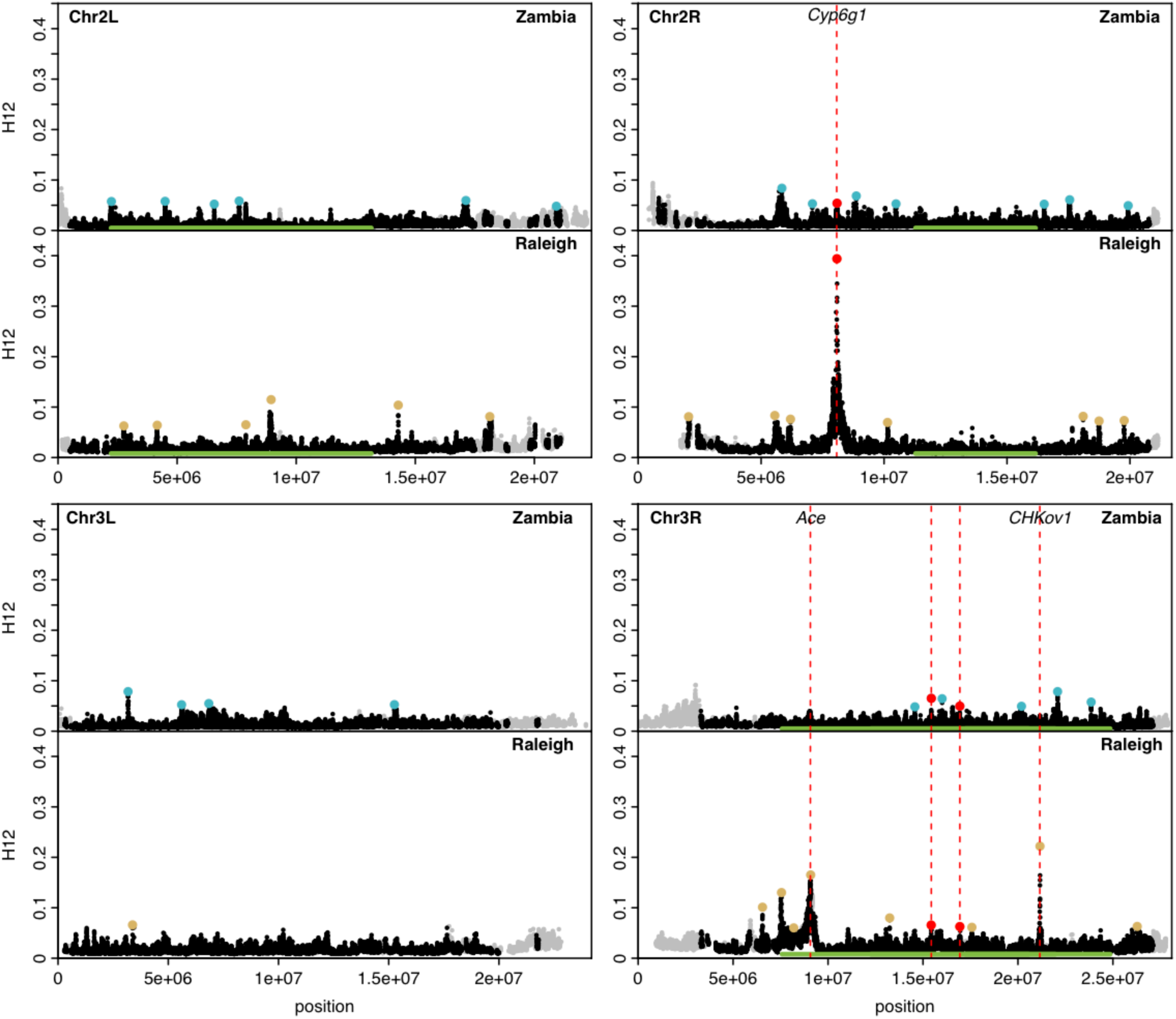
H12 scan in Zambian and Raleigh data. This figure is the same as Figure 2, except here the range of the y-axis is the same in both scans.

**Table S1:** Demographic parameters inferred by DaDi for models fit to Zambian data. Shown are parameter estimates and corresponding log likelihoods (LL) for the bottleneck, bottlegrowth, and growth models inferred by DaDi using Zambian short intron data Gutenkunst *et al*. 2009. *N_B_* represents the bottleneck size, *N_F_* the final population size, *T_B_* the duration of the bottleneck, and *T_F_* the duration of the final population size. For the growth model, both *N_F_* represents the final population size and *T* the time at which growth started. All population size estimates are in terms of units 4\**Ne*_ancestral_ and all time estimates are in terms of units 2\**Ne*_ancestral_.

**Bottleneck model:**
| $N_B$ | $N_F$ | $T_B$ | $T_F$ | LL |
| --- | --- | --- | --- | --- |
| 1.402 | 2.940 | 0.203 | 0.048 | -214.476 |

**Bottlegrowth model:**
| $N_B$ | $N_F$ | $T_B$ | $T_F$ | LL |
| --- | --- | --- | --- | --- |
| 1.465 | 3.388 | 0.128 | 0.116 | -214.376 |

| $N_F$ | $T$ | LL |
| --- | --- | --- |
| 2.87 | 0.17 | -228.54 |

**Table S2:** IDs of strains included in the analysis of LD and haplotype homozygosity patterns in Zambian and Raleigh data. We down sampled the Zambian and Raleigh strains used for the analysis of LD and haplotype homozygosity in Figures 1 and S1 to the 100 strains with the least amount of missing data for each population. This down sampling was done after extensive masking of regions of IBD, admixture, and inversions. Since the amount of missing data varied from chromosome to chromosome depending mainly on the presence of inversions, we down sampled 4 times for each population for each chromosome. The columns in this table show the strains that were included for the analysis for each chromosome for each population.

**Table S3:** Top 25 peaks in the Zambian (A) and Raleigh (B) data. Listed are the coordinates of the center of the analysis window with the highest H12 value in a peak, the edge coordinates of each peak, and the corresponding H12 and H2/H1 values in the analysis window.

**(A) Zambia**
| Chromosome | Start of peak | End of peak | Center of peak | H12 | H2/H1 |
| --- | --- | --- | --- | --- | --- |
| Chr2R | 5689984 | 5877052 | 5845467 | 0.084 | 0.448 |
| Chr3R | 22074333 | 22099126 | 22087580 | 0.079 | 0.106 |
| Chr3L | 3143696 | 3207496 | 3175369 | 0.079 | 0.106 |
| Chr2R | 8858953 | 8877924 | 8871146 | 0.069 | 0.210 |
| Chr3R | 15430749 | 15451721 | 15437579 | 0.065 | 0.591 |
| Chr3R | 15997772 | 16011492 | 16006285 | 0.064 | 0.132 |
| Chr2R | 17495878 | 17569226 | 17549291 | 0.061 | 0.151 |
| Chr2L | 16962865 | 17287118 | 17124919 | 0.060 | 0.368 |
| Chr2L | 7595364 | 7622463 | 7603947 | 0.058 | 0.156 |
| Chr2L | 4481025 | 4519970 | 4495822 | 0.058 | 0.208 |
| Chr3R | 23843073 | 23866972 | 23856074 | 0.058 | 0.143 |
| Chr2L | 2224927 | 2247036 | 2241538 | 0.058 | 0.125 |
| Chr3L | 6820734 | 6858547 | 6839441 | 0.055 | 0.141 |
| Chr2R | 8072574 | 8129240 | 8087869 | 0.054 | 0.345 |
| Chr2R | 7075343 | 7105539 | 7088009 | 0.053 | 0.407 |
| Chr2R | 10472210 | 10508206 | 10485863 | 0.053 | 0.449 |
| Chr3L | 15243946 | 15273825 | 15258107 | 0.053 | 0.592 |
| Chr3L | 5576197 | 5623343 | 5604719 | 0.052 | 0.222 |
| Chr2L | 6551349 | 6571793 | 6558631 | 0.052 | 0.194 |
| Chr2R | 16497602 | 16522034 | 16512033 | 0.052 | 0.274 |
| Chr3R | 16920082 | 16949847 | 16940157 | 0.050 | 0.266 |
| Chr3R | 20157200 | 20203615 | 20186573 | 0.049 | 0.262 |
| Chr2R | 19900056 | 19941093 | 19927955 | 0.049 | 0.167 |
| Chr3R | 14551505 | 14588594 | 14572745 | 0.048 | 0.622 |
| Chr2L | 20855660 | 21041236 | 20923022 | 0.048 | 0.671 |

**(B) Raleigh**
| Chromosome | Start of peak | End of peak | Center of peak | H12 | H2/H1 |
| --- | --- | --- | --- | --- | --- |
| Chr2R | 7723122 | 8452931 | 8082481 | 0.394 | 0.046 |
| Chr3R | 21100160 | 21196327 | 21161195 | 0.222 | 0.388 |
| Chr3R | 8561176 | 9347112 | 9084900 | 0.165 | 0.338 |
| Chr3R | 7441836 | 7656600 | 7549332 | 0.130 | 0.098 |
| Chr2L | 8787750 | 8981351 | 8946244 | 0.115 | 0.524 |
| Chr2L | 14260585 | 14297414 | 14284521 | 0.104 | 0.330 |
| Chr3R | 6503887 | 6568950 | 6548287 | 0.101 | 0.149 |
| Chr2R | 5488736 | 5706118 | 5556716 | 0.083 | 0.558 |
| Chr2R | 18048793 | 18139982 | 18096795 | 0.082 | 0.501 |
| Chr2L | 18066440 | 18156804 | 18128033 | 0.081 | 0.229 |
| Chr2R | 1980165 | 2741000 | 2051987 | 0.081 | 0.516 |
| Chr3R | 13235515 | 13272233 | 13244854 | 0.080 | 0.580 |
| Chr2R | 6158540 | 6220781 | 6194741 | 0.076 | 0.524 |
| Chr2R | 19741265 | 19784611 | 19765222 | 0.073 | 0.179 |
| Chr2R | 18730636 | 18755472 | 18743856 | 0.072 | 0.326 |
| Chr2R | 10116253 | 10172867 | 10141807 | 0.070 | 0.602 |
| Chr3L | 3361643 | 3387951 | 3379415 | 0.066 | 0.305 |
| Chr3R | 15417427 | 15448395 | 15431116 | 0.066 | 0.555 |
| Chr2L | 7874465 | 7898299 | 7886142 | 0.065 | 0.386 |
| Chr2L | 4146117 | 4176261 | 4164595 | 0.064 | 0.409 |
| Chr3R | 26262373 | 26301949 | 26281108 | 0.063 | 0.566 |
| Chr3R | 16908455 | 16953298 | 16939738 | 0.063 | 0.245 |
| Chr2L | 2735006 | 2781702 | 2762777 | 0.063 | 0.313 |
| Chr3R | 17548069 | 17614825 | 17573640 | 0.061 | 0.480 |
| Chr3R | 8113141 | 8551014 | 8193336 | 0.060 | 0.413 |

**Table S4:** Test for enrichment of peaks overlapping regions with inversions. We tested for an enrichment of peaks in regions of the genome with inversions that are present in at least 10 strains in the Zambian and Raleigh data sets. To do so we performed a one-sided binomial test calculating the probability of at least the number of peaks observed to overlap assuming a uniform distribution of peaks through the genome. We observe an enrichment of peaks overlapping common inversions on chromosome 3R when performing the scan in windows of 801 SNPs down sampled to 401 SNPs in Zambian data and in windows of 401 SNPs in Raleigh data (low probabilities are highlighted in bold). When we perform the scan in 801 SNP windows in Zambia, we find that the number of peaks overlapping this region decreases and is no longer significant, but instead we find an enrichment on chromosome 3L.

| Inversion | Number of overlapping peaks in Zambia (scan performed in 801 SNPs windows down sampled to 401 SNPs) | Probability | Number of overlapping peaks in Zambia (scan performed in 801 SNP windows) | Probability | Number of overlapping peaks in Raleigh (scan performed in 401 SNP windows) | Probability |
| --- | --- | --- | --- | --- | --- | --- |
| In(2L)t | 4 | 0.14 | 4 | 0.14 | 4 | 0.14 |
| In(2R)ns | 0 | 0.73 | 1 | 0.46 | 0 | 0.73 |
| In(3L)Ok | 3 | 0.20 | 5 | **0.02** | - | - |
| In(3R)K | 5 | 0.16 | 4 | 0.31 | 7 | **0.02** |
| In(3R)Mo | - | - | - | - | 2 | 0.31 |
| In(3R)P | 5 | **0.02** | 3 | 0.16 | 5 | **0.02** |

